# Genomic resources for the North American water vole (*Microtus richardsoni*) and the montane vole (*Microtus montanus*)

**DOI:** 10.1101/2021.04.04.438380

**Authors:** Drew J. Duckett, Jack Sullivan, Stacy Pirro, Bryan C. Carstens

## Abstract

**Background:** Voles of the genus *Microtus* are important research organisms, yet genomic resources in the genus are lacking. Providing such resources would benefit future studies of immunology, phylogeography, cryptic diversity, and more.

**Findings:** We sequenced and assembled nuclear genomes from two subspecies of water vole (*Microtus richardsoni*) and from the montane vole (*Microtus montanus*). The water vole genomes were sequenced with Illumina and 10X Chromium plus Illumina sequencing, resulting in assemblies with ~1,600,000 and ~30,000 scaffolds respectively. The montane vole was assembled into ~13,000 scaffolds using Illumina sequencing also. In addition to the nuclear assemblies, mitochondrial genome assemblies were also performed for both species. We conducted a structural and functional annotation for the best water vole nuclear genome, which resulted in ~24,500 annotated genes, with 83% of these receiving functional annotations. Finally, we find that assembly quality statistics for our nuclear assemblies fall within the range of genomes previously published in the genus *Microtus*, making the water vole and montane vole genomes useful additions to currently available genomic resources.

## Context

The genus *Microtus* consists of 62 species of voles distributed throughout North America, Europe, and Asia [1]. *Microtus* is believed to have experienced rapid speciation and diversification, with all speciation events occurring within the past four million years [2, 3], and it has been suggested that some nominal species, such as *M. pennsylvanicus*, contain cryptic diversity [4]. *Microtus* has been an important model system across multiple biological disciplines, including studies of adaptation (e.g., [5]), infectious disease (e.g., [6]), parental care (e.g., [7]), and population dynamics (reviewed in [8]). The rapid radiation of *Microtus* voles has hindered systematic classification, leading to multiple taxonomic revisions and conflicting phylogenetic analyses [1, 9, 10]. Consequently, both species boundaries and relationships among species are difficult to infer. Genomic resources within *Microtus* will help resolve these questions, and resources have steadily increased in recent years. Currently, four *Microtus* species have assembled genomes on GenBank, two European species (*M. agrestis* and *M. arvalis*) and two North American species (*M. ochrogaster;* [11], and *M. oeconomus*). The present study provides resources for two additional species: *M. richardsoni* and *M. montanus*.

The North American water vole (*M. richardsoni*) is adapted to a semiaquatic lifestyle, relying on alpine and sub-alpine streams for creating burrows and escaping predators [12]. Like other semiaquatic mammals (e.g., otters), it is likely that adaptations to this lifestyle have been driven by natural selection [13–15]. Water voles are among the largest species of *Microtus* and are known for making runways of stamped-down vegetation along streams through frequent movement [12, 16]. Unlike most other vole species, *M. richardsoni* does not appear to experience regular population boom and bust cycles, although population size in the species may be correlated to levels of precipitation [17]. Despite being listed as *Least Concern* by the IUCN Redlist [18], the species is listed as *Critically Imperiled* by the Wyoming Natural Diversity Database due to its specific habitat requirements, which can be substantially degraded by livestock grazing [19]. *Microtus richardsoni* occupies a large, disjunct distribution in the Pacific Northwest of North America, with habitat in the Cascades Mountains and the Rocky Mountains, spanning from southern Canada into central Utah. Four subspecies are currently recognized: *M. r. arvicoloides* in the Cascades Mountains, *M. r. richardsoni* in the Canadian Rocky Mountains, *M. r. macropus* in the central Rocky Mountains and Wyoming, and *M. r. myllodontus* in Utah. Due to the subspecific classifications and the disjunct range of the species, *M. richardsoni* has been included in multiple studies of phylogeography in the Pacific Northwest [20–22]. These studies were based solely on mitochondrial DNA, and the results of analyses that investigated species limits and demographic history were limited to inferences that can be derived from a single gene tree. Genomic resources for *M. richardsoni* will provide a rich source of data to address these knowledge gaps.

The montane vole (*M. montanus*) is partially sympatric with *M. richardsoni* and can be found throughout most of the water vole’s range with the exception of the Canadian Rockies. However, *M. montanus* can be found farther south and east including areas of California, Nevada, Colorado, Arizona and New Mexico [23]. The species has been divided into fifteen subspecies, including *M. m. canescens* in the Cascades Mountains., *M. m. nasus* in the central Rocky Mountains, and *M. m. amosus* in northern Utah. Notably, *M. montanus* does not exhibit a break in its range in the Columbia Basin, likely because it is not restricted to riparian areas like *M. richardsoni*. The species as a whole is listed as *Least Concern* by the IUCN Redlist, but *M. m. arizonicus* has been listed as endangered by the New Mexico State Game Commission Regulation [23], and *M. m. ricularis* has been noted as being of concern due to a small range and declining population size [24]. Genomic resources in *M. montanus* will provide a wealth of data to assess subspecies boundaries, quantify gene flow among subspecies, and aid in conservation efforts of threatened subspecies.

The present study provides two nuclear and one mitochondrial genome assembly for *M. richardsoni* along with single nuclear and mitochondrial genome assemblies for *M. montanus*. Furthermore, a structural and functional annotation are performed with one of the *M. richardsoni* genomes to aid in future studies of adaptation. Genome-level comparisons are made between the new genome assemblies and other *Microtus* genome assemblies to examine differences in assembly quality and repeat content.

### Sequencing and Nuclear Genome Assembly

Frozen tissue from a single *M. r. arvicoloides* individual collected from the southern Cascades Mountain range (JMS_292; 44.016667N, −121.750000E; [20]) was sent to Hudson Alpha (Huntsville, AL) for high molecular weight DNA extraction and 10X Chromium library preparation [25]. In the 10X method, each extracted DNA fragment receives a different barcode before the fragment is sheared for library preparation. After sequencing, these barcodes are used to connect sequencing reads for a more contiguous assembly. After sequencing with a single run on an Illumina HiSeqX, the resulting 150 base pair (bp) paired-end reads were input into Supernova for *de novo* genome assembly with --maxreads=all [26].

Additional tissue was obtained from a single *M. r. macropus* individual collected from the northern Rocky Mountains (JMG_88; 46.333333N, −114.633333E; [20]). DNA was extracted using a Qiagen DNeasy Blood and Tissue Kit, and the DNA was sent for library preparation and sequencing by Iridian Genomes, Inc (Bethesda, MD). 150bp paired-end reads were sequenced on two runs of an Illumina HiSeqX. Genome assembly was performed using two different deBruijn graph-based programs, SOAPdenovo and Discovar de novo [27, 28]. For SOAPdenovo, quality trimming was performed using fastQC and Trimmomatic with settings ILLUMINACLIP: 2:30:10, LEADING:3, TRAILING:3, SLIDINGWINDOW:4:15, and MINLEN:36 [29, 30]. SOAPdenovo assemblies were performed with settings max_rd_len=150, avg_ins=300, reverse_seq=0, asm_flags=3, rd_len_cutoff=150, rank=1, pair_num_cutoff=3, and map_len=32. SOAPdenovo was run with kmer values of 63, 89, 95, and 101 based on analysis of optimal kmer values in kmerGenie [31]. Raw reads were used as input for *de novo* genome assembly with Discovar as recommended in the program documentation.

In an attempt to provide the most contiguous assembly for *M. richardsoni*, a hybrid assembly was performed using the ARCS+LINKS pipeline [32, 33]. The ARCS+LINKS pipeline uses barcoding information from the 10X Chromium reads to scaffold the contigs from a separate genome assembly. Barcoded reads from *M. r. arvicoloides* were mapped to the *M. r. macropus* Discovar assembly with bwa mem [34] before converting the mapped reads to BAM format and sorting with SAMTools [35]. ARCS and LINKS were then run with settings –s 98 –c 5 –l 0 –z 500 –d 0 –r 0.05 –m 50-10000 –e 30000 and –d 4000 –k 20 –l 5 –t 2 –a 0.3 –o 0 –a 0.3 –z 500 respectively.

As part of a separate project, a single *M. montanus* individual from Utah (UMNH:Mamm:30891; 38.19381N, −111.5824E) was misidentified as *M. richardsoni*. DNA was extracted from the sample using a Qiagen DNeasy Blood and Tissue Kit before being sent to the University of California Davis Genome Center for library preparation and sequencing. Paired-end 150bp sequences were collected with a single shared run on an Illumina NovaSeq. Species identity was confirmed using the Barcode of Life Database (BOLD; [36]). Reads were checked and trimmed for quality with fastQC and Trimmomatic as above before mapping reads to the mitochondrial cytochrome oxidase I (COI) sequence of *M. r. macropus* [37] using bwa mem. The resulting mapped reads were converted to BAM format, sorted, and indexed with SAMTools. PCR duplicates were identified and removed with Picard [38], resulting reads were piled with SAMTools mpileup using base and mapping quality scores of 30, consensus sequences were generated with bcftools [39], and consensus sequences were converted to fastq format using vcfutils with a minimum depth filter of 5 and maximum depth filter of 10000 [35]. The resulting sequence was input into BOLD. Due to the low sequencing coverage, *de novo* genome assembly was not appropriate for *M. montanus*. To provide a preliminary genome sequence, a reference-guided genome assembly was performed with RaGOO [40]. Raw reads were input into Discovar to generate an initial genome assembly, misassembly correction was performed with RaGOO using reads trimmed with the same settings as the *M. r. macropus* reads, and RaGOO was then used to scaffold the Discovar contigs onto the *M. r. arvicoloides* assembly, which is more closely related to *M. montanus* than the other available *Microtus* genome assemblies [3]. Since *M. montanus* has less than half the chromosomes of *M. richardsoni* (2n = 22-24 in *montanus* versus 56 in *richardsoni* [41]), the possibility of structural errors in the *M. montanus* assembly was examined by calculating the percentage of reads that mapped back to the assembly using bwa mem and bamtools [42].

The final assemblies were submitted to GenBank [43], where screening was performed to identify any contamination, and contaminated scaffolds were removed. All assemblies were evaluated with QUAST [44], bbmap [45], custom Python scripts (https://github.com/djlduckett/Genome_Resources/), and BUSCO using the Euarchontoglires reference set [46]. After comparing assembly statistics from the different assemblies of *M. r. macropus*, the Discovar assembly was selected as best because it had less fragmentation, higher N50 and L50, and a higher BUSCO score than the SOAPdenovo assemblies (Table 1). Genome sequencing of *M. r. arvicoloides* produced over 800 million (M) reads and 47x genome sequencing coverage. The final genome assembly consisted of ~32 thousand (K) scaffolds with an N50 of 2.3 megabase pairs (Mb), 1.3% missing data (N), and a BUSCO score of 85.8%. Supernova estimated the length of the genome assembled to be ~2.4Gb and the total genome size to be ~2.6 gigabase pairs (Gb). *Microtus richardsoni macropus* sequencing produced over 600M reads and 35x coverage. Genome assembly with Discovar resulted in ~1.6M scaffolds with an N50 of 16 kilobase pairs (Kb), 0.06% Ns, and a BUSCO score of 54.5%. Given that there are many programs to perform *de novo* genome assembly from short reads, it is possible that another program would have produced a more contiguous *M. r. macropus* assembly, but previous studies have shown Discovar performs well compared to other programs [47, 48]. The hybrid assembly produced with the ARCS+LINKS pipeline had ~1.6M scaffolds, an N50 of 38Kb, 0.09% Ns, and a BUSCO score of 59.8%. Because of the poor quality of the hybrid assembly, it was not used for further analyses, and the *M. richardsoni* subspecies assemblies were kept separate. It seems likely that the high fragmentation of the hybrid assembly is due to the fragmentation of the Discovar input assembly. Published results with this hybrid pipeline often include a much higher sequencing coverage of the input contigs to produce a better starting point for the pipeline. Therefore, additional Illumina sequencing with *M. r. macropus* in the future could substantially improve the hybrid assembly. 108M reads (13x coverage) were used to produce the preliminary *M. montanus* genome, resulting in ~13K scaffolds, an N50 of ~3.1Mb, 8.8% Ns, and a BUSCO score of 82.6%. Additionally, 89.3% of reads mapped back to the *M. montanus* assembly.

**Table 1.**
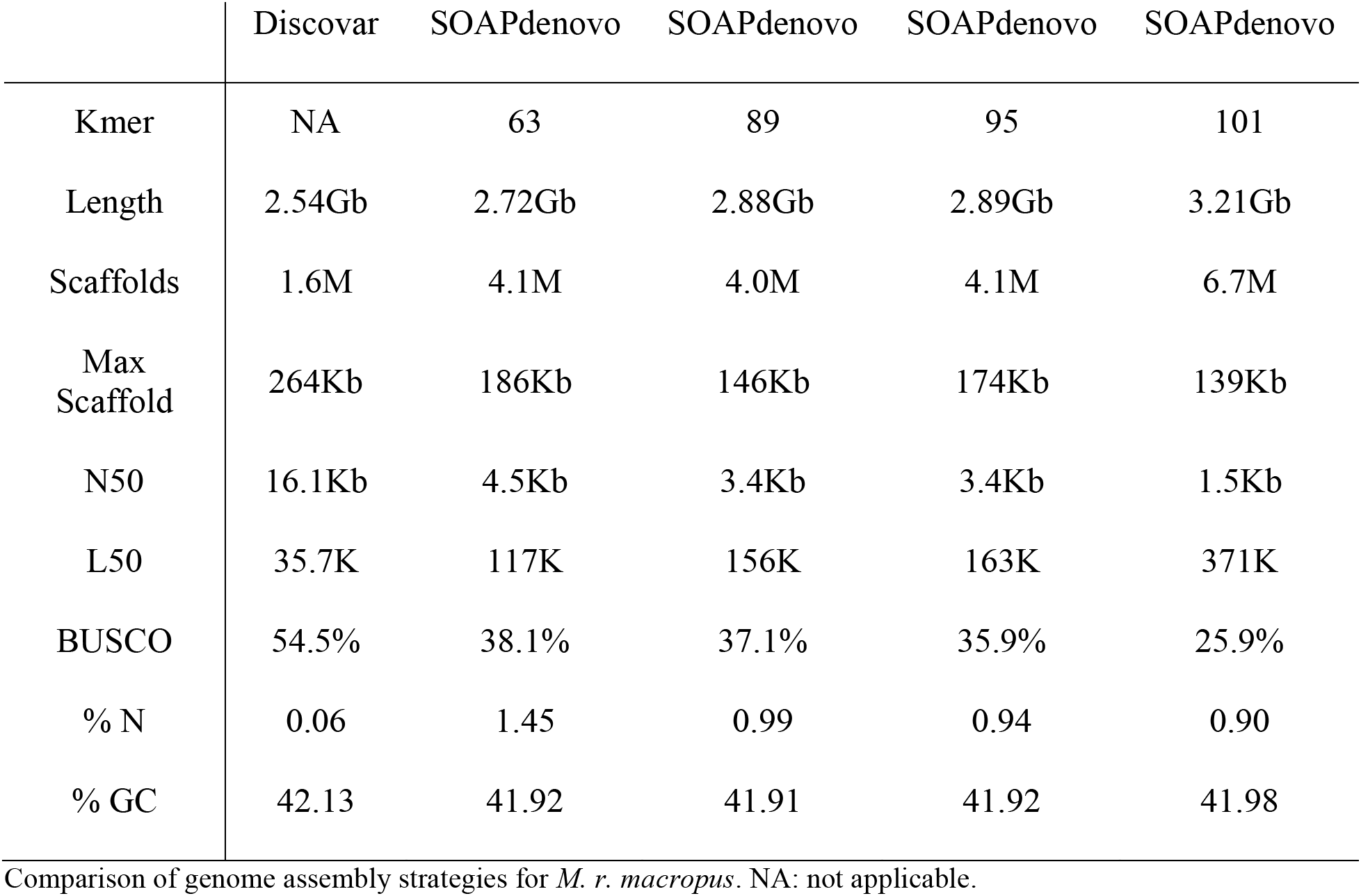

### Mitochondrial Genomes

The complete mitochondrial genomes of *M. r. arvicoloides* and *M. montanus* were assembled using the genomic sequencing reads. The mitochondrial genomes were assembled by both mapping reads to a reference mitochondrial genome and using the reference-guided assembly program Novoplasty [49]. For the mapping assembly, reads were mapped to the *M. r. macropus* mitochondrial genome, using the same steps as the *M. montanus* BOLD analysis. The mitochondrial assemblies were 16,285bp and 16,268bp in length with an average depth of coverage of 7886x and 6805x for *M. r. arvicoloides* and *M. montanus* respectively. Reference guided mitochondrial assemblies with Novoplasty used the *M. r. macropus* mitochondrial genome as the reference along with settings *Genome Range=12000-22000, K-mer=33, Read Length=150, and Insert size=400*. Because the *M. r. arvicoloides* dataset contained many reads, 25% of reads were subsampled to use for assembly, as suggested in the program documentation. The assemblies for *M. r. arvicoloides* and *M. montanus* were 16,298bp and 16,319bp in length with average depths of coverage of 5131x and 14,713x respectively. To compare mitochondrial assemblies between methods, the assemblies were aligned using the MUSCLE plugin in Geneious v. R9 with eight iterations and an open gap score of −1 [50, 51]. This comparison showed the Novoplasty assemblies contained multiple insertions compared to the mapped assemblies and the reference mitochondrial genome. These insertions were up to 13bp long in multiple genes, including trnT, trnK, and ATP8. Comparison to other *Microtus* mitochondrial genomes (*M. ochrogaster*; NC_027945.1 and *M. fortis*; NC_015243.1) showed that the Novoplasty assemblies were the only mitochondrial assemblies to exhibit these insertions. Therefore, the mapping assemblies were used for further analyses. The mapping assemblies for both species included ambiguous bases, which were much more frequent for *M. montanus* than *M. r. arvicoloides*. These may be the result of using the mitochondrial genome of a different subspecies (for *M. r. arvicoloides*) or species (for *M. montanus*) for mapping the reads. Additionally, the presence of nuclear DNA of mitochondrial origin (NUMTs; [52, 53]) may have influenced these results. If mitochondrial segments have been incorporated into the nuclear genomes and subsequent mutations have occurred, both nuclear and mitochondrial sequences could be mapped to the same mitochondrial region during assembly and result in the ambiguous bases observed here. It is likely that NUMTs are present, as they have been documented in other species of *Microtus* [54–56]. Both mitochondrial genomes were annotated using MITOS [57]. The annotations each consisted of 22 tRNA genes, 2 rRNA genes, and 13 protein coding genes.

### *Microtus* Genome Assembly Comparison

The available *Microtus* genome assemblies, *M. agrestis* (GCA_902806755.1), *M. arvalis* (GCA_007455615.1), *M. ochrogaster* (GCA_000317375.1), and *M. oeconomus* (GCA_007455595.1), were downloaded from GenBank. Assembly summary statistics were calculated using QUAST, bbmap, and custom Python scripts (https://github.com/djlduckett/Genome_Resources/). To compare repeat content among all genomes, including the three produced by the current study, repeats were first identified *de novo* using RepeatModeler [58]. RepeatMasker was then used to further identify repeats using a combined repeat library that included the repeats identified from RepeatModeler and those from the RepeatMasker *Rodentia* database [59]. The percentage of the genome consisting of each type of repeat element was extracted from the RepeatMasker log file for each genome assembly.

All genome assemblies used some form of Illumina sequencing (Table 2), although assembly continuity varied greatly among assemblies from 1366 scaffolds in *M. agrestis* to 1.6 M scaffolds in *M. r. macropus*. Genome coverage was similarly varied, from 13x in *M. montanus* to 35x in *M. r. macropus* to 77x in *M. arvalis* and *M. oeconomus*. The percent of repetitive regions ranged from 31.7% in *M. montanus* to 44.1% in *M. arvalis* (Figure 1), and repeat content did not appear to be associated with phylogenetic relatedness as repeats between the two subspecies of *M. richardsoni* were not more similar to each other than to other *Microtus* species. However, it is possible that the repeat content is affected by the continuity of the genome assemblies, and further research is needed to confirm this relationship.

**Table 2.**
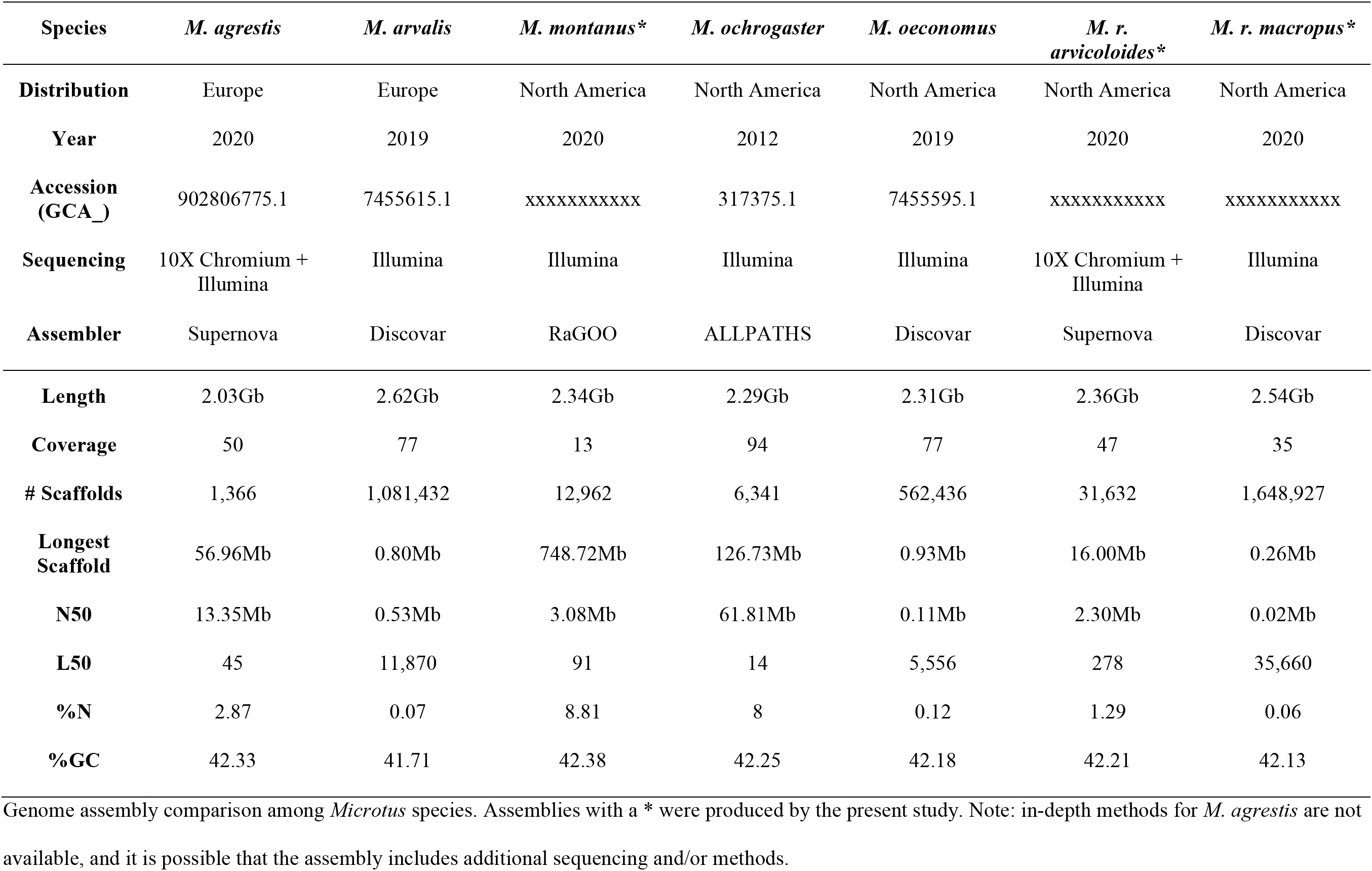

**Figure 1:**
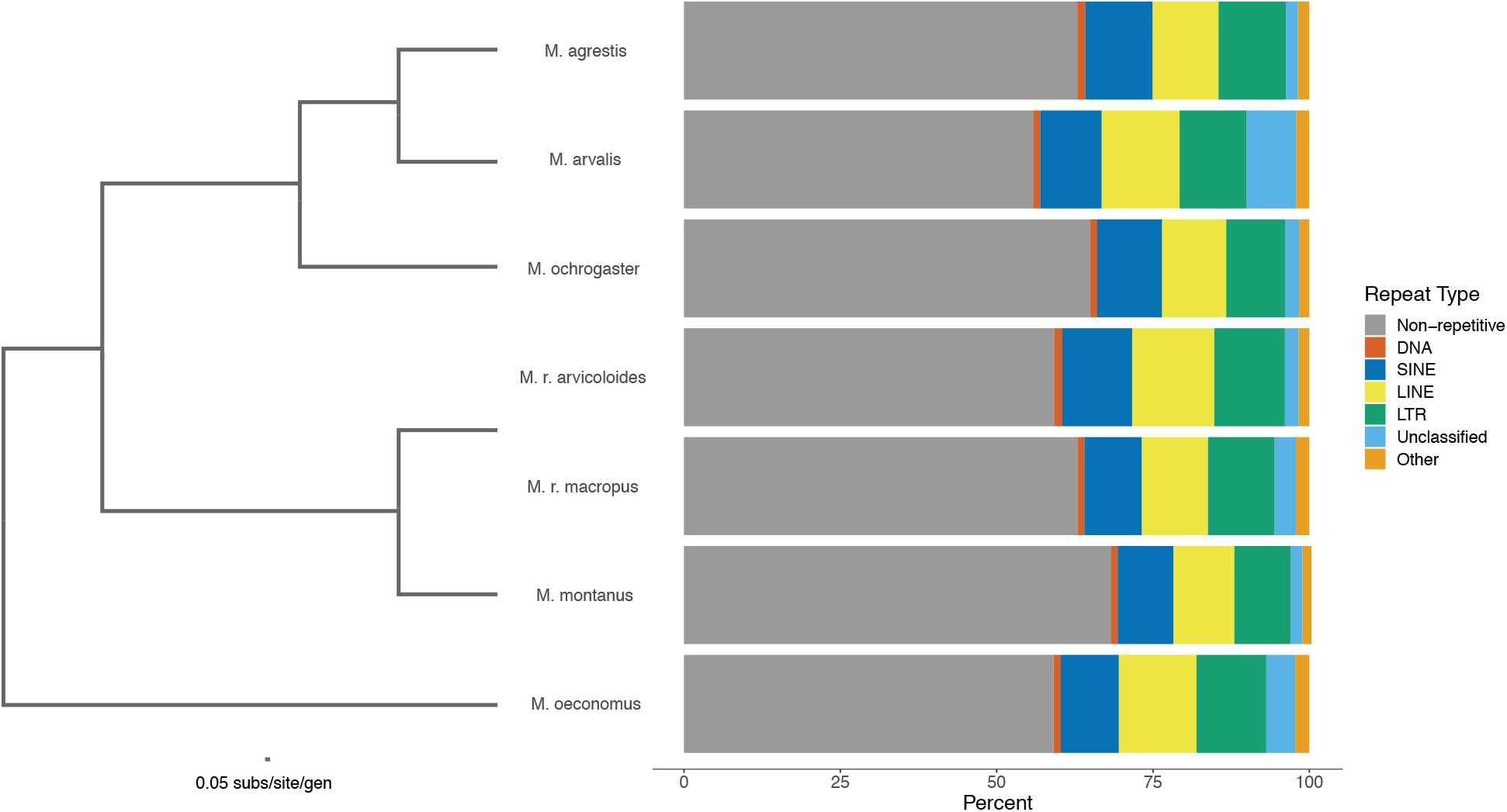
Repeat Content among *Microtus* genomes. SINE: short interspersed nuclear element; LINE: long interspersed nuclear element; LTR: long terminal repeat; Other consists of small RNA, satellite, simple, and low complexity repeats. The phylogeny displayed was recreated from [3] by pruning unincluded species from the data alignment and rerunning RAxML [75] with the same settings used in the original analysis.

### Genome Annotation

The *M. r. arvicoloides* genome assembly was annotated with the MAKER pipeline [60], loosely following the tutorial provided by Daren Card (https://gist.github.com/darencard/bb1001ac1532dd4225b030cf0cd61ce2). Briefly, the pipeline consists of masking repeats followed by multiple rounds of annotation with both evidence-based and ab-initio gene models. Repeats were identified as described above. Complex repeats were then extracted from RepeatMasker results using grep with keywords “Satellite” and “rich”. Within Maker, the model_org argument was set to “simple” so Maker would soft mask simple repeats, and the RepeatMasker results were provided to hard mask complex repeats. Evidence-based gene discovery used protein and mRNA sequences from the previous genome annotation of *M. ochrogaster* (GCF_000317375.1) as well as an additional RNASeq assembly from *M. pennsylvanicus* (GSM3499528; [61]). Hidden Markov models (HMMs) for ab-initio gene prediction were trained using both SNAP and Augustus [62, 63]. With SNAP, gene models identified by MAKER were filtered using an Annotation Edit Distance (AED) of 0.5 and an amino acid length of 50. After validating these models with SNAP’s Fathom utility, removing likely errors, and including 1000bp surrounding each training sequence, the training sequences were passed to the hmm-assembler script. For Augustus, training sequences plus 1000bp on each side were obtained from the first round of MAKER mRNA annotations. Augustus was used to train the HMM using the --long option in BUSCO and the Euarchontoglires reference set. MAKER was then run again with the previously annotated gene models and the HMM models from SNAP and Augustus. After the initial MAKER run, two cycles of ab-initio gene prediction and annotation with MAKER were performed. To prevent overfitting, results were compared after each round of MAKER. Because the increase in AED score was minimal between the first and second rounds of ab-initio gene prediction, further analysis was conducted on the results after the first round only. This round annotated ~24K genes with a mean gene length of 7445bp (Table 3), which is within the range found in previous studies of *M. ochrogaster* (22,427 genes; GCF_000317375.1) and *Arvicola amphibious* (25,136 genes; GCF_903992535.1). Of these annotations all occurred on scaffolds greater than 1Kb in length and 97% occurred on scaffolds greater than 10Kb in length.

**Table 3.**

|  | Before Gene Modeling | Gene Modeling Round 1 | Gene Modeling Round 2 |
| --- | --- | --- | --- |
| **Genes** | 20,945 | 24,548 | 23,811 |
| **Exons** | 139,845 | 192,974 | 179,225 |
| **mRNA** | 20,945 | 24,548 | 23,811 |
| **tRNA** | - | 24,504 | 24,539 |
| **5' UTR** | - | 1,229 | 1,180 |
| **3' UTR** | - | 503 | 642 |
| **Mean Gene Length** | - | 7,445 | 7,132 |
| **AED < 0.50** | 0.993 | 0.881 | 0.888 |
| **AED < 0.25** | 0.672 | 0.543 | 0.520 |
| **BUSCO (Complete)** | - | 67.7% | 70.5% |

Structural annotation summary after each round of MAKER. UTR: untranslated region; AED: annotation edit
distance. Values with dashes were not analyzed prior to gene modeling with SNAP and Augustus.

Functional annotation of the *M. r. arvicoloides* genome was performed using GOfeat, an online functional annotation tool that uses multiple protein databases including UniProt, InterPro, and Pfam [64–67]. An input file for GOfeat was generated by supplying the genome assembly FASTA file and the MAKER General Feature Format (GFF3) file to the Python package gffread [68]. GOfeat annotated 83.49% of genes. Biological Processes accounted for 42.46% of annotations, Cellular Components accounted for 30.29%, and Molecular Functions comprised 27.25%. The most frequent gene ontology (GO) terms were *positive regulation of transcription by RNA polymerase II, negative regulation of transcription by RNA polymerase II,* and *DNA-templated regulation of transcription* for Biological Processes, *cytoplasm* and *plasma membrane* for Cellular Components, and *metal ion binding* and *calcium ion binding* for Molecular Functions.

## Conclusion

The current study details the assembly and annotation of three nuclear and two mitochondrial genomes. Compared to previously published nuclear genomes, the *M. r. arvicoloides* and *M. montanus* genomes are of high quality as evidenced by the low number of scaffolds, high N50/L50 values, and high BUSCO scores. While not as complete as the other *Microtus* genomes, the nuclear genome of *M. r. macropus* will still be useful for mapping low coverage reads or reduced representation sequencing data. Furthermore, the mitochondrial genomes contributed here add to a growing number for the genus *Microtus* and reinforce earlier suggestions that high-quality mitochondrial genomes can be obtained as byproducts of nuclear sequencing (e.g., [69, 70]). Overall, the data presented serve as an example that even though they do not include chromosomal information, high-quality draft genomes can be produced from widely available and very cost-effective methods like the 10X Chromium protocol. These references can aid a variety of studies including those examining genus and species adaptation [71, 72], phylogenetics [10], phylogeography [22, 73], and disease dynamics [6, 74]. However, some activities, like exploring changes to chromosome structure, will not be possible due to the fragmentation and lack of chromosomal mapping for these assemblies. Finally, the *M. r. macropus* and *M. montanus* sequencing data and preliminary assemblies will serve as the building blocks of more accurate reference genomes in the future.

## Availability of Supporting Data and Materials

Raw sequences, nuclear assemblies, and mitochondrial assemblies are available from GenBank under BioProjects PRJNA673719, PRJNA509068, and PRJNA673873 for *M. r. arvicoloides*, *M. r. macropus*, and *M. montanus* respectively. The custom python script used to calculate genome assembly summary information is available on GitHub (https://github.com/djlduckett/Genome_Resources/). Full BUSCO tables, structural annotation gff files, functional annotation tables, and repeat libraries are available in the GigaScience data repository (http://gigadb.org/).

## Abbreviations

BOLD: Barcode of Life Database
bp: base pair
COI: cytochrome oxidase I
M: million
K: thousand
Gb: gigabase pairs
Mb: megabase pairs
Kb: kilobase pairs
SINE: short interspersed nuclear element
LINE: long interspersed nuclear element
LTR: long terminal repeat
AED: annotation edit distance
HMM: hidden Markov model
NUMT: nuclear DNA of mitochondrial origin

## Competing Interests

SP is the director of Iridian Genomes, Inc.

## Funding

Sequencing was funded by Iridian Genomes, Inc., as well as the National Science Foundation (DEB-1457519). Salary support for DD was provided by The Ohio State University and the National Science Foundation (DBI-1945347).

## Author Contributions

DD, JS, and BC conceived the study. JS, SP, and BC provided funding for sequencing. DD performed DNA extractions, assembled genomes, and annotated genomes with input from SP. DD and BC wrote the manuscript with input from JS and SP. DD and SP submitted the resources to GenBank.

## Acknowledgements

We thank Jeffrey Good and Eric Rickart/Utah Museum of Natural History for tissue samples, Michael Broe for advice with genome assembly and annotation, and the Ohio Supercomputer Center (OSC) for computational resources.

## Notes

https://www.ncbi.nlm.nih.gov/bioproject/673719

https://www.ncbi.nlm.nih.gov/bioproject/509068

https://www.ncbi.nlm.nih.gov/bioproject/673873

http://gigadb.org/

### Competing Interest Statement

Dr. Stacy Pirro is the director of Iridian Genomes, Inc.

